# SARS-CoV-2 harnesses host translational shutoff and autophagy to optimize virus yields: The role of the envelope (E) protein

**DOI:** 10.1101/2022.03.24.485734

**Authors:** Hope Waisner, Brandon Grieshaber, Rabina Saud, Wyatt Henke, Edward B Stephens, Maria Kalamvoki

## Abstract

The SARS-CoV-2 virion is composed of four structural proteins: spike (S), nucleocapsid (N), membrane (M), and envelope (E). E spans the membrane a single time and is the smallest, yet most enigmatic of the structural proteins. E is conserved among coronaviruses and has an essential role in virus-mediated pathogenesis. We found that ectopic expression of E had deleterious effects on the host cell as it activated stress responses, leading to phosphorylation of the translation initiation factor eIF2α and LC3 lipidation that resulted in host translational shutoff. During infection E is highly expressed although only a small fraction is incorporated into virions, suggesting that E activity is regulated and harnessed by the virus to its benefit. In support of this, we found that the γ_1_ 34.5 protein of herpes simplex virus 1 (HSV-1) prevented deleterious effects of E on the host cell and allowed for E protein accumulation. This observation prompted us to investigate whether other SARS-CoV-2 structural proteins regulate E. We found that the N and M proteins enabled E protein accumulation, whereas S prevented E accumulation. While γ_1_ 34.5 protein prevented deleterious effects of E on the host cells, it had a negative effect on SARS-CoV-2 replication. This negative effect of γ_1_ 34.5 was most likely associated with failure of SARS-CoV-2 to divert the translational machinery and with deregulation of autophagy pathways. Overall, our data suggest that SARS-CoV-2 causes stress responses and subjugates these pathways, including host protein synthesis (phosphorylated eIF2α) and autophagy, to support optimal virus production.

**Importance:** In 2020, a new β-coronavirus, SARS-CoV-2, entered the human population that has caused a pandemic resulting in 6 million deaths worldwide. Although closely related to SARS-CoV, the mechanisms of SARS-CoV-2 pathogenesis are not fully understood. We found that ectopic expression of the SARS-CoV-2 E protein had detrimental effects on the host cell, causing metabolic alterations including shutoff of protein synthesis and mobilization of cellular resources through autophagy activation. Co-expression of E with viral proteins known to subvert host antiviral responses such as autophagy and translational inhibition, either from SARS-CoV-2 or from heterologous viruses increased cell survival and E protein accumulation. However, such factors were found to negatively impact SARS-CoV-2 infection, as autophagy contributes to formation of viral membrane factories, and translational control offers an advantage for viral gene expression. Overall, SARS-CoV-2 has evolved mechanisms to harness host functions that are essential for virus replication.

## Introduction

Compared with other highly pathogenic coronaviruses (CoVs), the mortality rate of SARS-CoV-2 is approximately 2% among unvaccinated individuals (1). This mortality rate, along with the lack of pre-existing immunity, the fact that about 20% of infected individuals without pre-existing immunity require medical attention, and the highly transmissible nature of the virus, has led to the disruption of normal activities worldwide. Several highly effective vaccines have received use authorization, but the slow global vaccination rate and accumulation of adaptive mutations in different proteins of the virus, particularly the Spike protein, yield novel variants with different immunoevasion properties (2–5). Coronaviruses are enveloped viruses with a single-stranded, positive sense RNA genome. The coronavirus particle is composed of four structural proteins: nucleocapsid (N), membrane (M), envelope (E), and spike (S) (6). E is a small integral membrane protein that ranges from 75-106 aa (7–9). E protein localizes to the endoplasmic reticulum (ER), the ER-Golgi intermediate compartment (ERGIC) and the Golgi complex (10–15). The protein exists in different forms, including a monomeric form that potentially interacts with cellular proteins to alter the secretory machinery and to communicate signals, and a high-molecular weight homo-oligomer with a function in virion assembly (15–18). In addition, a pentameric form of E protein is an ion channel (viroporin) with mild selectivity for cations that has been linked to virus pathogenesis (19–29).

The importance of E during SARS-CoV and SARS-CoV-2 infections is highlighted by the fact that viruses lacking the gene for E protein display significantly reduced virus yields, due to aborted viral assembly that gives rise to immature virions with a strikingly aberrant morphology (12, 25, 30–32). For example, during infection with a mouse hepatitis virus (MHV) deleted of E, the virions display pinched, elongated rather than spherical shapes and smaller plaques, with irregular-shaped and jagged edges (33). How E protein facilitates virion morphogenesis remains unclear considering that only a small fraction of E is incorporated into the virions (34). A role of E in inducing membrane curvature has been proposed for MHV that is perhaps associated with E homo-polymerization and its interactors, but a mechanism is currently unknown (13).

The role for the cation channel activity of E during SARS-CoV-2 infection is also unclear although mutations within the transmembrane domain that inhibit the ion channel activity in SARS-CoV E are reversed by this virus (35). However, as most known mutations that impair the ion channel activity of E also impair E oligomerization, it is currently unknown if one or both properties of the protein are rescued (21, 25, 36–38). The transport of Ca^2+^ by SARS-CoV E has been correlated with inflammatory-mediated lung damage *in vivo*, highlighting the importance of E in viral pathogenesis (39–45). The channel activity of E could also alter the secretory pathway or the luminal environment, leading to efficient trafficking of virions (18, 27, 46, 47). Consistently, some of the proposed interactors of E are associated with ion transport and others with vacuoles and mitochondria, suggesting that E may participate in re-organizing membranes and the recruitment of lipid processing machineries at sites of virion assembly (11, 48–54).

Considering that E protein localizes in the ER-ERGIC-Golgi compartments and forms an ion channel we sought to determine the type of responses activated in cells ectopically expressing E. We found that E protein triggered ER-signaling pathways that led to phosphorylation of the translation initiation factor eIF-2α with a concomitant translational shutoff and LC3 lipidation. Both effects indicate that major metabolic alterations occur in cells expressing E that impact protein synthesis and potentially mobilize energy resources. We also found that E protein accumulation was restricted in cells ectopically expressing E protein. To further understand the functions of E we determined whether proteins from heterologous viruses known to prevent eIF-2α phosphorylation and LC3 lipidation could reverse the adverse effects of E on the host. The γ_1_ 34.5 protein of HSV-1 is known to prevent host translational shutoff during HSV-1 infection by recruiting the protein phosphatase 1α (PP1α) to dephosphorylate eIF-2α, which is phosphorylated by activated protein kinase R (PKR) following foreign RNA sensing (55–59). In addition, γ_1_ 34.5 protein inhibits autophagy by binding to the autophagy-inducing protein Beclin-1 that is downstream of activated PKR (59–62). Mutant HSV-1 viruses lacking the Beclin-1-interacting domain of γ_1_ 34.5 display reduced viral replication *in vitro* and *in vivo*, due to robust activation of autophagy (59–66). We found that γ_1_ 34.5 could reverse eIF-2α phosphorylation, but not LC3 lipidation induced by E, and enabled E protein accumulation.

An interesting observation was that HSV-1 γ_1_ 34.5 inhibited SARS-CoV-2 infection. One mechanism was through inhibition of the host translational shutoff by γ_1_ 34.5 that is imposed by the virus to gain translational advantage over the host. Additionally, disruption of autophagy pathways by γ_1_ 34.5 during SARS-CoV-2 infection led to formation of aberrant vacuolar structures, most likely containing engulfed organelles, instead of viral membrane factories. Taken together, our data suggest that SARS-CoV-2 harnesses stress response pathways of the host for optimal progeny virus production.

## Results

### The E protein of SARS-CoV-2 initiates autophagy and interferes with translation initiation

The E protein accumulates in the ER and ERGIC where it can form a channel with weak cation specificity which may exhibit Ca^2+^ transport activity in the ERGIC (10, 11, 14, 15) (19-22, 24-26, 28). While only a small amount of E protein expressed during SARS-CoV-2 infection is incorporated into the virions, the protein appears to also induce membrane curvature, and participate in membrane scission (13, 34, 67–69). Thus, we sought to determine ER signaling responses that may be activated by E expression. We found that ectopic expression of E in HEK-293 cells caused LC3 lipidation that was apparent by 48 h post-transfection (Figure 1A). Moreover, we observed reduced p62/SQSTM1 accumulation (Figure 1B). The p62/SQSTM1 is an adaptor protein that sorts ubiquitinated cargo to autophagosomes for degradation and subsequently is degraded itself (70, 71). This reduction of p62/SQSTM1 accumulation is indicative of autophagy activation. We did not observe changes in the levels of optineurin (OPTN) suggesting that mitophagy was not induced by E expression (Figure 1B). Also, we did not observe changes in the levels of ATG5 protein (autophagy related 5), which along with ATG12 protein acts as an E1-activating enzyme during autophagy (Figure 1B) (72). In addition, we tested whether E expression could trigger phosphorylation of the translation initiation factor eIF-2α, a modification that is usually observed when unfolded protein response (UPR) pathways are activated (73, 74). We observed that ectopic expression of E triggered accumulation of p-eIF-2α (Figure 1C). As a control, HEK-293 cells were infected with an HSV-1 γ_1_ 34.5-null mutant, which cannot reverse phosphorylation of eIF-2α. Finally, following infection of Caco-2 cells with SARS-CoV-2 we observed some accumulation of p-eIF-2α (Figure 1D). We conclude that ectopic expression of E protein activates stress responses that lead to phosphorylation of eIF-2α and LC3 lipidation.

**Figure 1:**
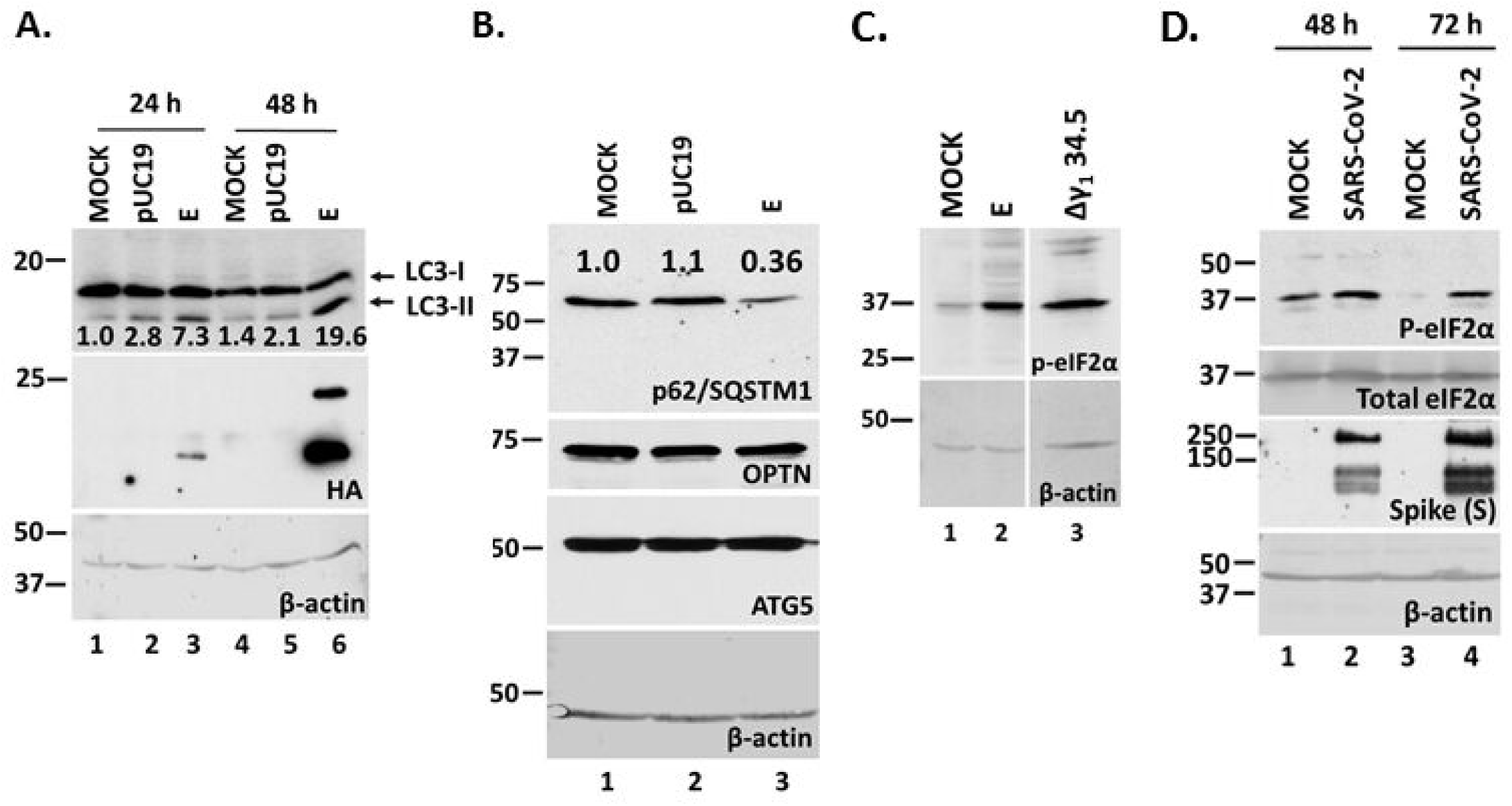
SARS-CoV-2 E expression causes LC3 lipidation, reduction in p62/SQSTM1 levels, and increased eIF-2α phosphorylation. **A:** HEK-293 cells were either left untransfected, transfected with a control pUC19 plasmid, or an E-HA expressing plasmid. The cells were harvested at 24 and 48 h post-transfection and equal amounts of proteins were analyzed for LC3 lipidation and E expression. The ratio of LC3-II/LC3-I is shown below. **B:** Transfections were as in panel A. Equal amounts of proteins were analyzed for p62/SQSTM1, Optineurin and ATG5. **C:** HEK-293 cells were transfected with an E-HA expressing plasmid or infected with a HSV-1 Δγ_1_ 34.5 virus (5 PFU/cell). The cells were harvested at 48 h post-transfection or at 14 h post-infection. Equal amounts of proteins were analyzed for p-eIF-2α. **D:** Caco-2 cells were infected with SARS-CoV-2 (2 PFU/cell). The cells were harvested at 48 h and at 72 h post-infection and equal amounts of proteins were analyzed for p-eIF-2α and total eIF-2α. Spike served as a control for the infection. β-actin served as a loading control in panels A-D.

### The γ_1_ 34.5 protein of HSV-1 inhibited phosphorylation of eIF-2α but not LC3 lipidation induced by E expression

The γ_1_ 34.5 protein of HSV-1 is known to prevent host translational shutoff (55–58, 63) and autophagy (60). To assess if γ_1_ 34.5 could reverse the effects of E protein, HEK-293 (Figure 2A) or A549 (Figure 2B) cells were co-transfected with an E and a γ_1_ 34.5-expressing plasmid. Cells co-transfected with the E-expressing plasmid and an empty vector served as a control. Additional controls included cells transfected with the individual plasmids and untransfected cells. The cells were harvested at 48 h post-transfection and equal amounts of proteins were analyzed for p-eIF-2α. As shown in Figures 2A-B, E expression triggered accumulation of p-eIF-2α (see also manuscript by Henke W. et al) that was blocked by the presence of γ_1_ 34.5. The phosphorylation of eIF-2α due to E expression caused translational shutoff (Figure 2C, lanes 3-4) that was partially reversed in the presence of γ_1_ 34.5 (Figure 2C, lane 5). In a similar transfection assay, we tested if γ_1_ 34.5 could inhibit LC3 lipidation triggered by E expression. As shown in Figure 2D, LC3 lipidation due to E expression was not reversed by γ_1_ 34.5. This is perhaps because γ_1_ 34.5 interferes with phagophore elongation through Beclin-1 binding, which does not necessarily interfere with LC3 lipidation. We conclude that γ_1_ 34.5 protein inhibits the phosphorylation of eIF-2α triggered by E expression, but does not inhibit LC3 lipidation.

**Figure 2:**
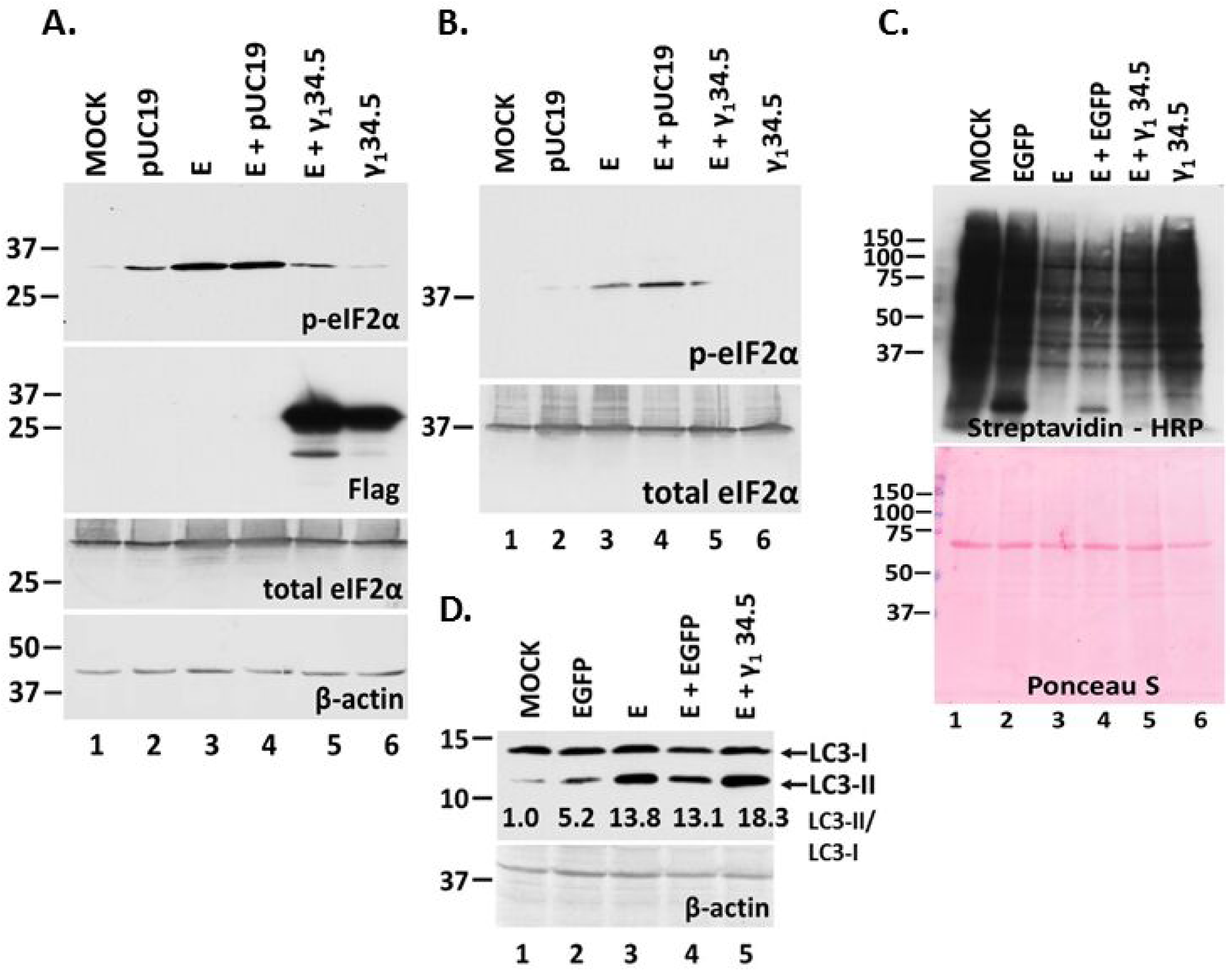
The γ_1_ 34.5 protein of HSV-1 prevents p-eIF-2α accumulation induced by E expression but not LC3 lipidation. **A:** HEK-293 cells were either left untransfected, or transfected with the control plasmid pUC19, an E-HA expressing plasmid, co-transfected with E-HA and pUC19, co-transfected with E-HA and γ_1_ 34.5-expressing plasmids, or with a γ_1_ 34.5-expressing plasmid (Flag-tagged). The cells were harvested at 48 h post-transfection and equal amounts of proteins were analyzed for p-eIF-2α, total eIF-2α, or γ_1_ 34.5 protein expression (Flag tagged). β-actin served as a loading control. **B:** Experiment was as in panel A, but it was performed in A549 cells. **C:** HEK-293 cells were either left untransfected, or transfected with the control plasmid pLenti CMV GFP Puro expressing EGFP, an E-HA expressing plasmid, co-transfected with E-HA and pLenti CMV GFP Puro, co-transfected with E-HA and γ_1_ 34.5-expressing plasmids, or with a γ_1_ 34.5-expressing plasmid (Flag-tagged). At 46 h post-infection the cells were starved for 3 h in RPMI Medium 1640 without L-methionine and subsequently incubated with medium supplemented with Click-iT AHA reagent for 2 h. Click chemistry reaction to monitor nascent protein synthesis was performed as described in the Materials and Methods. Both a ponceau S staining of the membrane and the reaction of HRP with the ECL substrate (Pierce) are depicted. **D:** Transfections in HEK-293 cells and protein analysis were as in panel A and analyzed for LC3 with the ratio of LC3-II to LC3-I shown below. The pUC19 plasmid was replaced with pLenti CMV GFP Puro that expresses the control protein EGFP. β-actin served as a loading control.

### The γ_1_ 34.5 protein of HSV-1 and the SARS-CoV-2 M and N proteins allowed for E protein accumulation

Both p-eIF-2α and LC3 lipidation could prevent E protein accumulation. To test this, we determined if γ_1_ 34.5 protein expression impacted E expression. HEK-293 cells were co-transfected with vectors expressing E and γ_1_ 34.5. Cells co-transfected with the E-expressing plasmid and a plasmid expressing EGFP, or cells transfected with the individual plasmids served as controls. The cells were harvested at 48 h post-transfection and the accumulation of E protein was assessed by analyzing equal amounts of cell lysates by western blot. The levels of E protein were lower when co-expressed with GFP compared with E alone, most likely because of competition of the two plasmids for transport into the nucleus, gene transcription and protein translation. However, when E was co-expressed with the γ_1_ 34.5 protein, accumulation of E protein was strongly enhanced (Figures 3A). In contrast, we did not observe N protein accumulation when it was co-expressed with the γ_1_ 34.5 protein (Figure 3B). These data suggest that stress responses activated following E expression negatively impact E accumulation, however, HSV-1 γ_1_ 34.5 protein can reverse these effects.

**Figure 3:**
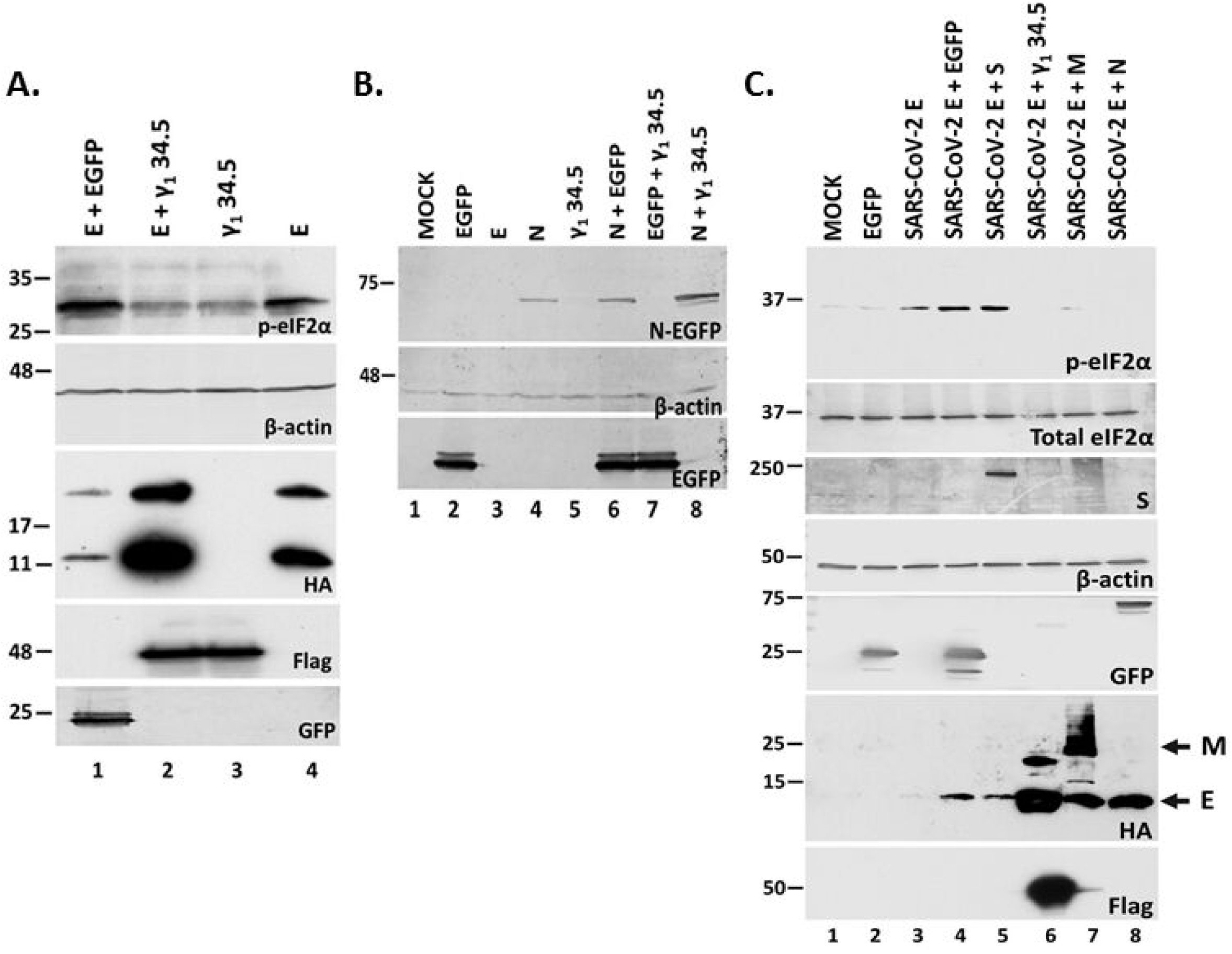
The γ_1_ 34.5 protein of HSV-1 and the SARS-CoV-2 M and N proteins resulted in E protein accumulation. **A.** HEK-293 cells were transfected with an E-HA expressing plasmid, a γ_1_ 34.5-expressing plasmid (Flag-tagged), co-transfected with E-HA and γ_1_ 34.5-expressing plasmids, or E-HA and EGFP-expressing plasmids. The cells were harvested at 48 h post-transfection and equal amounts of proteins were analyzed for p-eIF-2α, E-HA, γ_1_ 34.5 expression (Flag tagged), or GFP. β-actin served as a loading control. **B.** Transfections were as in panel A. Cell lysates prepared at 48 h post-transfection were analyzed using a GFP antibody with β-actin serving as a loading control. **C.** HEK-293 cells were transfected with an E-HA expressing plasmid, an EGFP-expressing plasmid, or co-transfected with an E-HA expressing plasmid and plasmids expressing SARS-CoV-2 M, N, S, or the HSV-1 γ_1_ 34.5 protein, respectively. Single transfections were done using 500 ng per well and co-transfections were performed using 1 μg per well (500 ng per plasmid). The cells were harvested at 24 h post-transfection and equal amounts of proteins were analyzed for expression of E-HA, S-HA, M-HA, γ_1_ 34.5 (Flag tagged), or GFP (control protein GFP and N fused to GFP). β-actin served as a loading control. Arrows indicate the E and M proteins that are both tagged with an HA epitope.

Considering that signaling responses activated following E expression have deleterious effects on the host, SARS-CoV-2 must control E functions to ensure optimal virus replication (39–45). Thus, we sought to determine if any SARS-CoV-2 proteins can reverse E effects allowing for E protein accumulation. We chose to analyze the effects of other virion proteins that may impact E functions through interactions. HEK-293 cells were co-transfected with plasmids expressing the E protein and either the M, N, or S proteins of SARS-CoV-2. HEK-293 cells were also co-transfected with plasmids expressing the E protein and either the EGFP or the HSV-1 γ_1_ 34.5 protein to serve as negative and positive controls, respectively. As shown in Figure 3C, both N and M proteins prevented p-eIF-2α accumulation due to E expression and enhanced E protein accumulation. In contrast, the S protein did not prevent p-eIF-2α accumulation and did not support E protein accumulation. Nevertheless, the γ_1_ 34.5 protein was more effective than the M and N proteins of SARS-CoV-2 in enabling E protein accumulation. We also assessed the impact of different proteolytic machineries on E protein accumulation and found no significant effect (supplemental data and Figure S1). Thus, E protein expression negatively impacts its own accumulation but this effect is reversed by proteins from SARS-CoV-2 or by heterologous viruses that appear to counterbalance E effects.

### SARS-CoV-2 E homologs and E oligomerization mutants trigger phosphorylation of eIF-2α

In the next series of experiments, we determined whether specific E mutants could decrease the propensity of E protein to induce ER stress responses. Two point mutations were inserted in the transmembrane domain of the E protein, asparagine (N) at position 15 was converted to alanine (A) and the valine (V) at position 25 was converted to phenylalanine (F) (N15A/V25F). These mutations are known to reduce E oligomerization to some extent (21, 22, 36, 75). The N15A mutation reduces pentamerization of E, while V25F reduces higher order oligomers (75). As shown in Figure 4A, the E N15A/V25F-expressing cells accrued similar levels of p-eIF-2α as cells expressing wild-type E (compare lane 4 to lane 3), suggesting that reduced E oligomerization does not reduce ER stress responses triggered by E. This level of p-eIF-2α was again reversed by γ_1_ 34.5 protein (compare lane 8 to lane 4). We also tested a mutant of E in which the conserved proline at position 54 was changed to a glycine (E-P54G). P54 is located within the cytoplasmic domain at the turn of a β-coil-β motif and likely affects E topology. E-P54G also triggered p-eIF-2α that was partially reversed by γ_1_ 34.5 protein (compare lane 5 to lane 3, and lane 9 to lane 5). LC3 lipidation was triggered by the unmodified E protein and all mutants tested, albeit to a greater extent by E-P54G. We conclude that disruption of E pentamerization or oligomerization does not reduce ER stress responses triggered by E expression.

**Figure 4:**
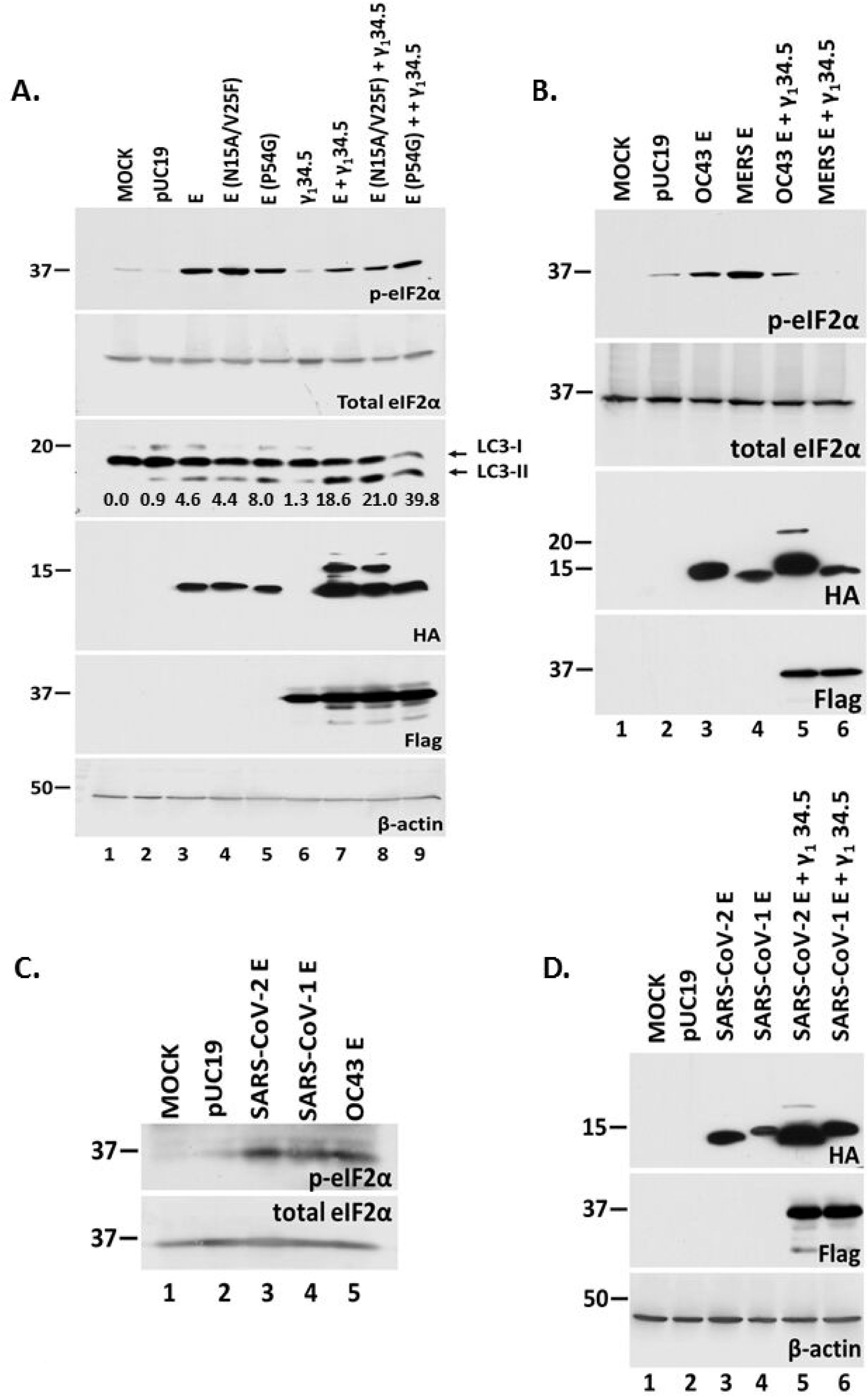
Homologs of E or oligomerization mutants do not abrogate phosphorylation of eIF-2α. **A.** HEK-293 cells were transfected with plasmids expressing either wild-type E or various E oligomerization mutants. In addition, HEK-293 cells were transfected with a γ_1_ 34.5-expressing plasmid, or co-transfected with various E forms and a γ_1_ 34.5-expressing plasmid. The cells were harvested at 48 h post-transfection and equal amounts of proteins were analyzed for p-eIF-2α, total eIF-2α, E-HA, LC3, or γ_1_ 34.5 expression (Flag tagged). The ratio of LC3-II to LC3-I is also shown. β-actin served as a loading control. **B, C, D.** Plasmids encoding for E protein from different CoVs including SARS-CoV, MERS-CoV, and HCoV-OC43 were transfected in HEK-293 cells or co-transfected with a γ_1_ 34.5-expressing plasmid. The cells were harvested at 48 h post-transfection and equal amounts of proteins were analyzed for p-eIF-2α, total eIF-2a, E-HA, and γ_1_ 34.5 expression (Flag-tagged). β-actin served as a loading control.

We then sought to determine whether the E protein of SARS-CoV, MERS-CoV, and HCoV-OC43 trigger similar responses as SARS-CoV-2 E (31). Cells were co-transfected with E-expressing and γ_1_ 34.5 -expressing plasmids as described above. Like SARS-CoV-2 E protein, SARS-CoV, MERS-CoV, and HCoV-OC43 E homologs induced phosphorylation of eIF-2α (Figure 4B-C). In each case the HSV-1 γ_1_ 34.5 protein blocked phosphorylation of eIF-2α (Figure 4B) and enabled E protein accumulation (Figure 4B and D). We conclude that E homologs and E oligomerization mutants display the same propensity as SARS-CoV-2 E to induce ER stress responses that result in eIF-2α phosphorylation.

### The γ_1_ 34.5 protein of HSV-1 reverses the translational shutoff imposed by SARS-CoV-2 and restricts virus infection

Considering that HSV-1 γ_1_ 34.5 protein could antagonize E functions important for virus replication such as translational shutoff, we determined the effect of γ_1_ 34.5 on SARS-CoV-2 infection. We developed a Vero E6 cell line expressing γ_1_ 34.5 protein under a tetracycline inducible promoter (Tet ON) from an integrated lentiviral vector. At 48 h after inducing γ_1_ 34.5 expression, the Vero E6 + γ_1_ 34.5 cell line was infected with the reporter virus icSARS-CoV-2-mNG (0.0001 PFU/cell) and GFP expression was compared with infected Vero E6 cells. Expression of HSV-1 γ_1_ 34.5 protein resulted in fewer cells that were positive for GFP compared with the control cells (Figure 5A). Expression of γ_1_ 34.5 protein at 48 h following induction with doxycycline was confirmed (Figure 5B). We also found that infection with either the wild-type virus (Figure 6A) or the reporter virus (Figure 6B) triggered phosphorylation of eIF-2α in Vero E6 cells, but not in γ_1_ 34.5 -expressing cells.

**Figure 5:**
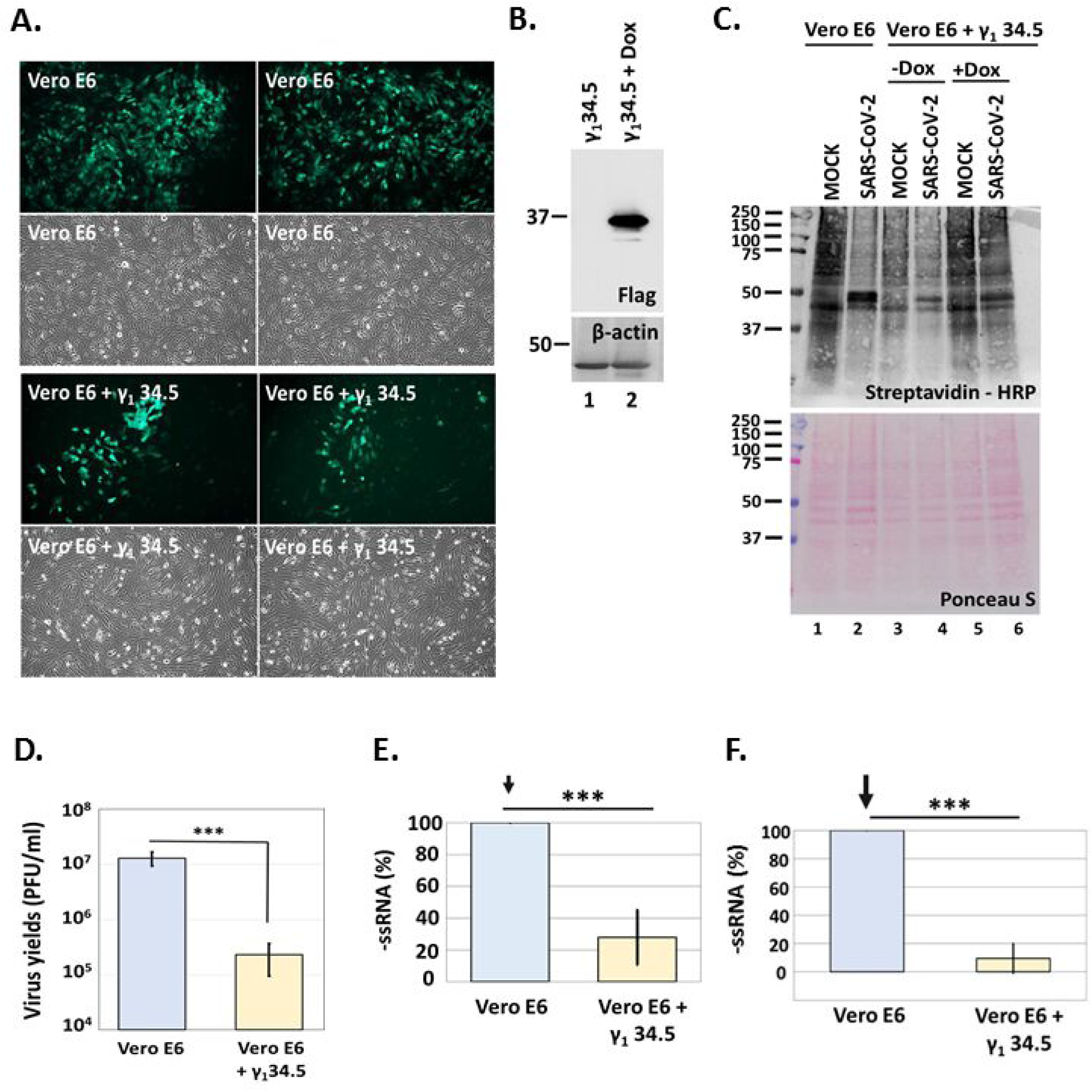
HSV-1 γ_1_ 34.5 inhibits SARS-CoV-2 infection. **A.** Vero E6 + γ_1_ 34.5 cells were treated with doxycycline (5 μg/ml for 48 h) to induce γ_1_ 34.5 expression. Induced cells along with parental Vero E6 cells were infected with icSARS-CoV-2-mNG (10^-4^ PFU/cell). Images were captured at 24 h post-infection using an Olympus microscope. **B.** Expression of γ_1_ 34.5-Flag protein following doxycycline treatment (20 μg/ml) of Vero E6 + γ_1_ 34.5 cells for 48 h. **C.** Vero E6 and Vero E6 + γ_1_ 34.5, either untreated or treated with doxycycline (20 μg/ml) to induce γ_1_ 34.5 expression, were infected with SARS-CoV-2 (10^-4^ PFU/cell). At 34 h post-infection the cells were starved for 3 h in RPMI Medium 1640 without L-methionine (Thermo-Fisher) and subsequently incubated with medium supplemented with Click-iT AHA (L-azidohomoalanine) reagent (Invitrogen) for 2 h. Cells were lysed in a solution containing 1% SDS in 50 mM Tris-HCl, pH 8.0, and labelled proteins were reacted with biotin-alkyne (PEG4 carboxamide-propargyl biotin) in a Click-chemistry reaction according to manufacturer’s instructions using the Click-iT™ Protein Reaction Buffer Kit (Invitrogen). Biotinylated proteins were analyzed in a denaturing polyacrylamide gel and detected with streptavidin-HRP. Both a ponceau S staining of the membranes and the reaction of HRP with 4-chloro-1-naphthol supplemented with hydrogen peroxide are depicted. **D.** Infections were performed with SARS-CoV-2 (10^-4^ PFU/cell) in replicate cultures of Vero E6 or doxycycline-treated (20 μg/ml) Vero E6 + γ_1_ 34.5 cells. The cells were harvested at 24 h post-infection and intracellular progeny virus was quantified by a plaque assay in Vero E6 cells. **E-F.** Infections were performed with either the wild type (panel E) or the reporter virus (panel F) as in panel D, in replicate cultures. Cells were harvested at 24 h post-infection and the –ssRNA was quantified by real time PCR analysis.

**Figure 6:**
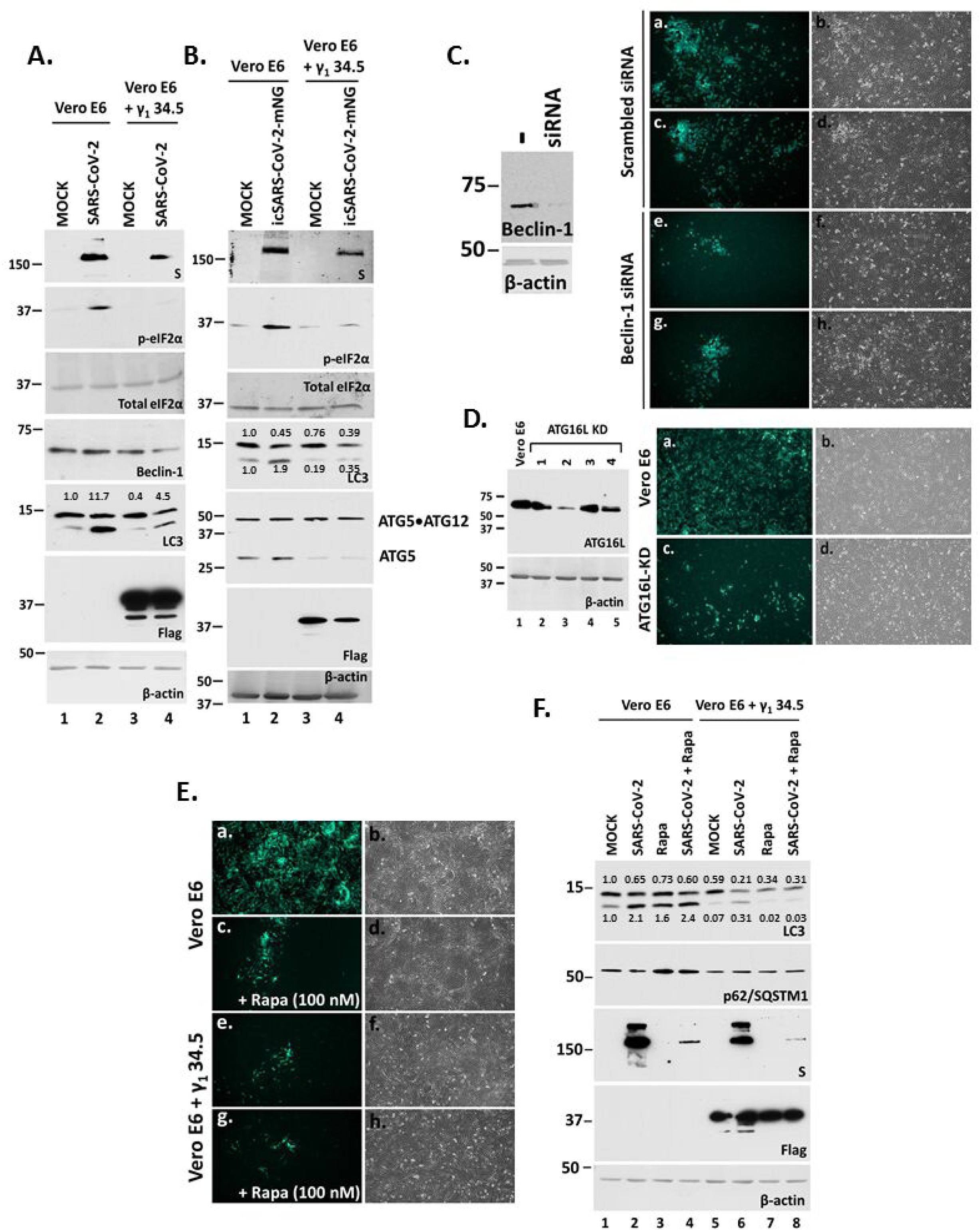
Effect of γ_1_ 34.5 protein on autophagy during SARS-CoV-2 infection. **A-B.** Replicate cultures of Vero E6 and doxycycline-induced Vero E6 + γ_1_ 34.5 cells were infected with either the SARS-CoV-2 USA-WA1/2020 (panel A) or the reporter virus icSARS-CoV-2-mNG (panel B) (10^-4^ PFU/cell). Cells were harvested at 36 h post-infection and equal amounts of proteins were analyzed for p-eIF-2α, total eIF-2α, LC3, γ_1_ 34.5 (Flag-tagged), ATG5, Beclin-1, S protein expression, and β-actin. Numbers represent ratio of LC3-II/LC3-I (panel A) and quantification of LC3-I and LC3-II (panel B). **C.** Vero-E6 cells were transfected at 50% confluency with either a control (scrambled) siRNA (Santa Cruz; sc-37007) or Beclin 1 siRNA (Santa Cruz; sc-29797) using Lipofectamine 3000 according to manufacturer’s instructions (Invitrogen). Both siRNAs were used at a 300 nM concentration and the cells were transfected for 72 h before infection. Efficiency of Beclin-1 depletion is depicted. Vero E6 cells treated with either the scrambled siRNA, or the Beclin-1 siRNA as above were infected with icSARS-CoV-2-mNG (10^-4^ PFU/cell). Images were captured at 24 h post-infection using an Olympus microscope. **D.** Vero E6 and ATG16L KD derivatives were infected with icSARS-CoV-2-mNG, as above and images were captured at 24 h post-infection. Efficiency of ATG16L depletion using a specific shRNA (Sigma) expressed from an integrated lentiviral vector is depicted. Four different cell lines were established using four different shRNAs (1–4) to deplete ATG16L and the cell line expressing shRNA #2 was selected as depletion of ATG16L was more efficient. **E.** Vero E6 + γ_1_ 34.5 cells were treated with doxycycline for 48 h to induce γ_1_ 34.5 expression, as above. Induced cells along with parental Vero E6 cells were infected with icSARS-CoV-2-mNG (10^-4^ PFU/cell). Rapamycin was added to the cultures together with the virus at 100 nM. Images were captured at 24 h post-infection using an Olympus microscope. **F.** Vero E6 + γ_1_ 34.5 cells that were treated with doxycycline (20 μg/ml) to induce γ_1_ 34.5 expression as above, along with Vero E6 cells were infected with SARS-CoV-2 (10^-4^ PFU/cell). Rapamycin was added to the cultures together with the virus at 10 μM. Cells were harvested at 36 h post-infection and equal amounts of proteins were analyzed for LC3 lipidation, p62/SQSTM1, and S protein. Quantification of LC3-I and LC3-II is depicted. Expression of γ_1_ 34.5 was verified with a Flag antibody. β-actin served as a loading control.

We then asked if phosphorylation of eIF-2α leads to host translational shutoff during SARS-CoV-2 infection, and whether it could be prevented by γ_1_ 34.5 protein expression. The Vero E6 + γ_1_ 34.5 cell line that was either not induced or induced to express γ_1_ 34.5 protein and parental cells were infected with SARS-CoV-2 (0.0001 PFU/cell). Cells were starved of methionine for 3 h starting at 34 h post-infection followed by labelling with the amino acid analog of methionine L-azidohomoalanine. Click-chemistry was used to detect the labelled proteins. As shown in Figure 5C, SARS-CoV-2 infection caused translational shutoff (lane 2 compared with lane 1) that was prevented in the presence of γ_1_ 34.5 (lane 6 compare with lane 2). An intermediate phenotype was observed in infected, doxycycline-untreated Vero E6 + γ_1_ 34.5 cells (lane 4 compare with lanes 2 and 6), perhaps due to some leakiness of γ_1_ 34.5 protein expression in un-induced cells. Staining with ponceau S of total proteins served as a loading control.

To quantify the effect of γ_1_ 34.5 on SARS-CoV-2 infection, the Vero E6 + γ_1_ 34.5 cell line that was induced to express γ_1_ 34.5 protein and Vero E6 cells were infected with SARS-CoV-2 (0.0001 PFU/cell) and progeny virus production was quantified at 24 h post-infection using plaque assays. The presence of γ_1_ 34.5 protein caused an approximate 100-fold reduction in infectious virus production at 24 h post-infection (Figure 5D). In a similar assay, we compared the amounts of the negative-strand RNA (-ssRNA) of the virus in the γ_1_ 34.5 – expressing cell line versus parental cells. We found that the presence of γ_1_ 34.5 protein caused a 70% and a 90% reduction in the –ssRNA of the WT-virus and the reporter virus, respectively (Figure 5E-F). Overall, γ_1_ 34.5 was able to inhibit the translational shutoff imposed during SARS-CoV-2 infection and which resulted in a decrease in virus replication and progeny virus production.

### γ_1_ 34.5 protein alters autophagic responses during SARS-CoV-2 infection

During SARS-CoV-2 infection autophagy appears to supplement the viral membrane factories with membranes and metabolites required for virus replication (76–78). While γ_1_ 34.5 is known to interfere with autophagy by binding to Beclin-1, we observed that co-expression of γ_1_ 34.5 protein with E did not reduce, but rather enhanced LC3 lipidation (Figure 2D). LC3 lipidation was also induced during SARS-CoV-2 infection, however in the presence of γ_1_ 34.5, the levels of both non-lipidated and lipidated LC3 were reduced after infection with either the wild type virus (panel 6A, compare lane 4 to lane 2) or the reporter virus (panel 6B, compare lane 4 with lane 2). We also noticed a decrease in the amounts of S protein, Beclin-1 and ATG5 in infected γ_1_ 34.5 - expressing cells, although the levels of the ATG5/ATG12 complex remained unaltered. These data indicated that γ_1_ 34.5 disrupted autophagic responses during SARS-CoV-2 infection, causing a reduction in viral infection.

The γ_1_ 34.5 protein is known to combat autophagy during HSV-1 infection through both a direct mechanism, by interacting with Beclin-1, and an indirect mechanism, by inhibiting PKR-induced phosphorylation of eIF-2α. To assess the impact of Beclin-1 on SARS-CoV-2 infection we depleted cells of Beclin-1 using a specific siRNA followed by infection with the reporter virus. Depletion of Beclin-1 caused a delay in SARS-CoV-2 infection (Figure 6C). Additionally, depletion of ATG16L, a critical factor for synthesis of the autophagosome precursor, reduced SARS-CoV-2 infection (Figure 6D). These data suggest that early autophagy events are essential during SARS-CoV-2 infection.

While early autophagy events are critical during SARS-CoV-2 infection, treatment with the autophagy inducer rapamycin was inhibitory, most likely due to degradation of factors critical for virus infection or virions *per se* (Figure 6E). The rapamycin effect could not be reversed by γ_1_ 34.5 (Figure 6E). A striking observation was that γ_1_ 34.5 sensitized the cells and the levels of both lipidated and non-lipidated LC3 and the levels of the autophagy adaptor p62/SQSTM1 were reduced following exposure to an autophagy inducer, including SARS-CoV-2 or rapamycin (Figure 6F). Notably, depletion of Beclin-1 during SARS-CoV-2 infection increased LC3 lipidation (data not shown).

Finally, we determined if γ_1_ 34.5 expression could cause overt changes in the viral membrane factories through its effects on autophagy. For this, doxycycline-treated Vero E6 + γ_1_ 34.5 cells and parental cells that were either exposed to SARS-CoV-2 or not, were processed for TEM analysis. Extensive membrane rearrangements and aberrant vesicular structures were observed in the cytoplasm of SARS-CoV-2 infected cells compared with uninfected cells (Figure 7, compare panels b-d to panel a). However, infection of γ_1_ 34.5 –expressing cells resulted in formation of oversized vacuoles that contained what appeared to be trapped cellular organelles, including mitochondria and endosomes undergoing degradation (Figure 7, compare panels f-h to panel e). A potential engulfment or fusion event with an organelle resembling a lysosome has been marked with a red arrow (Figure 7, panel h). We conclude that γ_1_ 34.5 expression during SARS-CoV-2 infection altered autophagic responses and caused formation of abnormal vacuoles with different organelles entrapped undergoing degradation.

**Figure 7:**
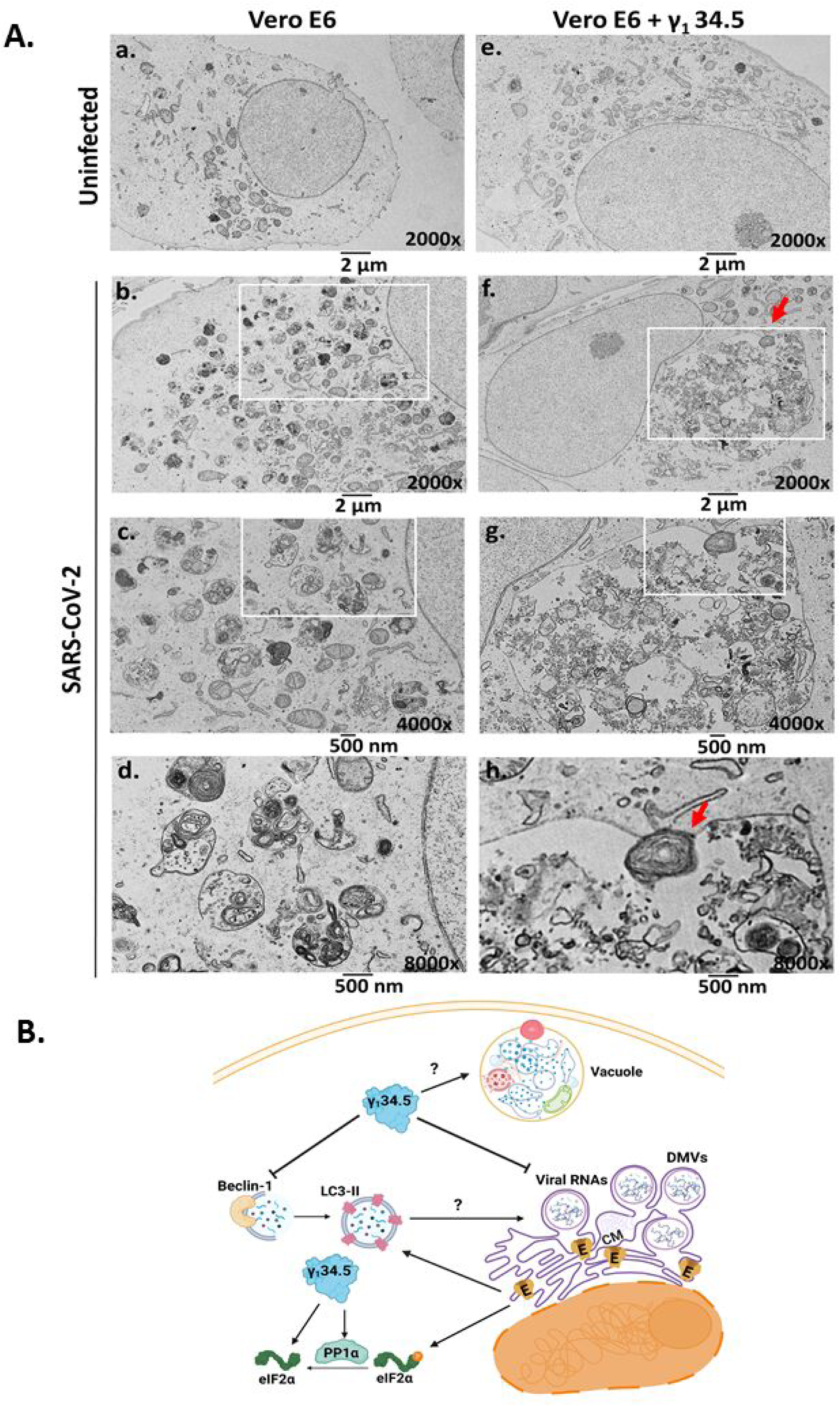
Aberrant vacuolar structures in the cytoplasm of SARS-CoV-2 infected γ_1_ 34.5 – expressing cells. **A.** Replicate cultures of Vero E6 and doxycycline-induced Vero E6 + γ_1_ 34.5 cells seeded on coverslips were infected with SARS-CoV-2 USA-WA1/2020 (10^-4^ PFU/cell). The cells were fixed with 2% glutaraldehyde at 42 h post-infection and processed for TEM analysis, as detailed in materials and methods. At least 50 cells were analyzed per sample. Representative images are depicted. **B.** Model summarizing the mechanism of interference of SARS-CoV-2 replication by the HSV-1 γ_1_ 34.5 protein. SARS-CoV-2 infection causes extensive reorganization of the ER/ERGIC compartments that leads to formation of viral membrane factories where the virus replicates. It also imposes a translational shutoff that offers an advantage to viral over host genes for expression, while suppressing host defense gene expression. E protein could contribute to virus replication by facilitating membrane rearrangements through activation of autophagy that supports the growth of the viral factories. E along with other viral proteins could be responsible for the translational shutoff during SARS-CoV-2 infection. γ_1_ 34.5 protein could disrupt autophagy activated during SARS-CoV-2 infection by binding to Beclin-1 and/or through other mechanisms. Also, γ_1_ 34.5 protein suppresses host translational shutoff during SARS-CoV-2 infection. Both effects could result in inhibition of SARS-CoV-2 replication. Autophagic vacuoles with abnormal size and morphology observed in SARS-CoV-2 infected γ_1_ 34.5 -expressing cells could represent defects in the autophagolysosome pathway.

## Discussion

Our studies emanated from the observation that SARS-CoV-2 causes major rearrangements in the ER-Golgi membranes that form the viral membrane factories where the virus replicates and virions assemble. Since E is a small transmembrane protein that oligomerizes in the ER and the ERGIC, we hypothesized that it could disrupt the functions of these organelles (10–12, 14). Indeed, we observed that E protein caused LC3 lipidation, a hallmark of autophagy initiation, and phosphorylation of the translation initiation factor eIF-2α resulting in host translational shutoff. It is likely that E expression activates ER-stress responses, including the unfolded protein response that results in PERK activation, which phosphorylates eIF-2α (73, 74, 79). Disruption of ER homeostasis could subsequently lead to autophagy activation.

Coronaviruses are known to impose host translational shutoff during the early stages of infection to prevent the infected host from synthesizing new proteins, while translation of viral mRNAs is not affected (80–93). Phosphorylation of eIF-2α has been reported during infection by SARS-CoV, SARS-CoV-2, the transmissible gastroenteritis virus (TGEV) and other CoVs and both kinases PERK and PKR appear to participate in this process (94–100). PERK could be activated following disruption of ER homeostasis by viral proteins accumulating in the ER (101, 102). For example, E could disrupt ER Ca^2+^ homeostasis through its viroporin function. PKR could be activated following viral RNA sensing. Ectopic expression of several SARS-CoV proteins including S and ORF3a triggers p-eIF-2α due to disruption of ER homeostasis (103–106).

We observed that ectopic expression of E protein had deleterious effects on the host cell that impacted E protein accumulation. It is not uncommon for viruses to develop mechanisms to regulate and harness the activity of proteins that trigger deleterious responses to ensure optimal replication. To test this, we co-expressed E with either other SARS-CoV-2 proteins that are known to interact with E, or with proteins from heterologous viruses that can evade host translational shutoff and autophagy. We discovered that the γ_1_ 34.5 protein of HSV-1 inhibited phosphorylation of eIF-2α triggered by E expression and allowed for E protein accumulation. The mechanism by which γ_1_ 34.5 protein prevents accumulation of p-eIF-2α has been previously described. During HSV-1 infection, PKR is activated and phosphorylates eIF-2α. HSV-1 γ_1_ 34.5 recruits the PP1α phosphatase to dephosphorylate eIF-2α (56, 57, 61, 62). While γ_1_ 34.5 protein inhibited accumulation of p-eIF-2α in E-expressing cells, it did not inhibit LC3 lipidation. γ_1_ 34.5 protein can inhibit autophagy by binding to the phagophore nucleation factor Beclin-1 and by inhibiting different pattern recognition receptors and downstream effectors like the TANK-binding kinase 1 (TBK1) (60, 107, 108). Recent studies demonstrated that depletion of Beclin-1 had little effect on LC3 lipidation, but it played a critical role during autophagosome formation and macromolecule degradation through the autophagy pathway (109). These observations provide an explanation why LC3 lipidation occurs when E protein is co-expressed with γ_1_ 34.5.

Besides γ_1_ 34.5, we found that the M and N proteins of SARS-CoV-2 prevent accumulation of p-eIF-2α in E-expressing cells, while S exacerbated accumulation of p-eIF-2α. Both M and N interact with E during virion assembly (6). It is likely that this binding alters the localization of E, its oligomerization status, or its potential binding with host factors, which alters its propensity to cause ER stress responses. Alternatively, the immunoevasion properties of M and N could reverse the translation inhibition imposed by E (110–112). On the other hand, ectopic expression of S is known to trigger phosphorylation of eIF-2α, and in the presence of E such an effect was exacerbated (103). Consistent with this, we have observed increased LC3 lipidation when E and S were co-expressed ectopically (data not shown).

The properties and functions of the E proteins are generally conserved among beta coronaviruses, as the E proteins of SARS-CoV, MERS-CoV, and HCoV-OC43 were found to trigger similar responses as SARS-CoV-2 E (31). Mutations that obstructed the ion channel function of E or decreased E oligomerization did not suppress p-eIF-2α accumulation. Perhaps the intracellular localization of E and its interactors are sufficient to trigger ER stress responses leading to eIF-2α phosphorylation.

An interesting observation was that SARS-CoV-2 displayed decreased replication and progeny virus production in γ_1_ 34.5-expressing cells (Figure 7B). One explanation is that by preventing host translational shutoff, γ_1_ 34.5 decreased the efficiency of viral gene expression, and enabled expression of host defense genes that are known to combat SARS-CoV-2 infection (80, 113). An additional possibility was that γ_1_ 34.5 disrupted autophagy pathways utilized by SARS-CoV-2. Coronaviruses are known to exploit autophagosome formation to support DMV biogenesis, while stalling lysosome fusion to evade autophagy-mediated degradation. Transient depletion of Beclin-1, which functions as a scaffold in forming a multiprotein assembly during autophagy initiation and nucleation, and is a known target of γ_1_ 34.5, obstructed the infection. In addition, depletion of ATG16L, an integral part of the complex involved in LC3 lipidation that is essential for autophagosome formation and expansion, had a negative effect on SARS-CoV-2 infection. Thus, it appears that LC3 lipidation is essential for SARS-CoV-2 DMV formation. Another striking observation was that infection of γ_1_ 34.5-expressing cells with SARS-CoV-2 led to formation of enormous size vacuoles, almost half the size of the nucleus, containing what appeared to be various organelles undergoing degradation. This abnormal phenotype of vacuoles may be the result of either defective autophagy or defective proteolysis. While SARS-CoV-2 through ORF3a can inhibit fusion of autophagosomes with lysosomes and decrease lysosomal activity by increasing lysosomal pH, this was not sufficient to yield the abnormal vacuole phenotype, and γ_1_ 34.5 was required to sensitize the cells by altering early autophagy events (114).

Overall, we provide novel evidence that ectopic expression of E causes adverse effects on the host cell. These effects were found to be antagonized by γ_1_ 34.5, a protein from a heterologous virus. This led us to discover that the activity of E is likely regulated during SARS-CoV-2 infection by other viral proteins to ensure optimal virus production. Finally, we demonstrated that pathways inhibited by γ_1_ 34.5 are required for optimal SARS-CoV-2 growth, therefore these pathways could be considered novel antiviral targets.

## Materials and Methods

### Cells and Viruses

The A549 cells (human lung adenocarcinoma), Caco-2 (human colorectal adenocarcinoma), HEK-293 (human embryonic kidney epithelial cells), and Vero E6 (normal monkey kidney epithelial cells) were obtained through ATCC. The SARS-related coronavirus 2 isolate USA-WA1/2020 was obtained through BEI resources (NR-52281). The icSARS-CoV-2-mNG was obtained through the World Reference Center for Emerging Viruses and Arboviruses (“WRCEVA”) at the University of Texas Medical Branch at Galveston (“UTMB”).

### Development of a Vero E6 cell line expressing γ_1_ 34.5 under tetracycline inducible promoter from an integrated lentiviral vector

A plasmid expressing the γ_1_34.5 ORF with a FLAG-tag was digested with Hind III and Xba I to extract only the FLAG-tagged γ_1_34.5. This fragment was then inserted into the pLenti-mCherry-Mango II x 24 plasmid (Addgene #127587) digested with Nhe I and BamH I. HEK-293 cells were seeded in a 60 mm dish at 60% confluence and were co-transfected with the pLenti-mCherry-FLAG-γ_1_34.5 plasmid described above, the Gag-Pol-expressing plasmid, and the vesicular stomatitis virus G (VSV-G)-expressing plasmid at a ratio of 7:7:1 (5 μg total amount of DNA) using Lipofectamine 3000 (Invitrogen) according to the manufacturer’s instructions. At 48 h after transfection, the supernatant from the cultures was collected, filtered through a 0.45-μm-pore-size filter, and used to infect Vero-E6 cells, as described before (115, 116). Puromycin selection initiated at 24 h after exposure of cells to lentiviruses and continued until only resistant clones survived. Resistant cultures were then plated in a 6-well plate and exposed to doxycycline (5 μg/ml) for 48 h. After 48 h the cells were harvested in triple lysis solution and equal amounts of lysates were analyzed for expression of FLAG-γ_1_34.5 via immunoblot analysis. Cultures with the greatest expression of FLAG-γ_1_34.5 were then used for all further experiments.

### Plasmids and transfection assays

The genes for the E proteins were synthesized by SynBio Technologies with HA-tags fused to the C-terminus of each protein. The genes were expressed in the pcDNA3.1 (+) vector. The vectors expressing the SARS-CoV-2 M and S proteins (with C-terminal HA-tag) were obtained from Sino Biologicals. The pcDNA3.1 (+)-N-eGFP-N plasmid, expressing N from SARS-CoV-2, was obtained from GenScript Biotech (Catalog # MC_0101137).

Transfections of HEK-293 cells, seeded in 12-well plates, were performed using Lipofectamine 3000 according to the manufacturer’s instructions. Unless stated otherwise, all transfections were done using 1 μg of DNA total for single transfections, or 1.5 μg total for co-transfections (750 ng per plasmid). Cells were harvested at 48 h post-transfection and equal amounts of protein were analyzed by immunoblot analysis.

### Western blot analyses

The procedures for immunoblotting were described elsewhere (see also supplemental materials and methods) (115, 117).

### Monitoring of nascent protein synthesis

Cells were uninfected or infected with SARS-CoV-2 for indicated times. Starvation of cells was done for 3 h by incubating with RPMI Medium 1640 without L-methionine (Thermo-Fisher). Cells were then incubated with the same medium supplemented with the Click-IT™-AHA (L-Azidohomoalanine) reagent (Invitrogen) for 2 h. Cells were lysed in a solution containing 1% SDS in 50 mM Tris-HCl, pH 8.0. Click-chemistry reaction for protein detection was performed using biotin alkyne (PEG4 carboxamide-propargyl biotin) according to manufacturer’s instructions using the Click-iT™ Protein Reaction Buffer Kit (Invitrogen). Labeled proteins were separated by SDS-PAGE. The membrane containing the biotin alkyne labeled proteins was incubated in a solution containing 1% BSA with streptavadin-HRP (Invitrogen) for subsequent visualization.

### Detection of viral negative sense –ssRNA

Cell lysates were collected in TRIzol reagent (Ambion) at indicated times post-infection. Total RNA was extracted via phenol-chloroform extraction method. Reverse-transcription PCR was then performed using LunaScript RT Master Mix Kit (NEB) using a gene specific primer. This primer was used to specifically detect the negative strand RNA from SARS-CoV-2, as described previously (118). The reverse transcription primer, 5’- ACAGCACCCTAGCTTGGTAGCCGAACAACTGGACTTTATTGA -3’, contains IAC (internal amplification control) tag-2 and a part targeting the ORF1ab gene of SARS-CoV-2. Real-time PCR was then performed using SYBR Green reagent (Invitrogen) according to the manufacturer’s recommendations in a 7500 fast real-time PCR system (Applied Biosystems). The forward primer, 5’- AGGTGTCTGCAATTCATAGC-3’ (743-762bp), and the reverse primer, 5’- ACAGCACCCTAGCTTGGTAG -3’ (IAC tag-2), were used for amplification.

### Processing cells for transmission electron microscopy (TEM)

Cell monolayers on Thermanox plastic coverslips (13 mm) (Nunc) were fixed with 2% glutaraldehyde in 0.1 M sodium cacodylate buffer, pH 7.4, and washed two times with 0.1 M sodium cacodylate buffer. Samples were post-fixed in 1% osmium tetroxide plus 1.5% potassium ferrocyanide in 0.1 M sodium cacodylate for 30 min at room temp and rinsed 3 times with distilled water. Samples were dehydrated in a graded series of ethanol as follows: 50%, 70%, 80%, 95%, 100%, 100%. A drop of Embed 812 resin was applied to each coverslip and samples embedded on Thompson molds were polymerized overnight at 60°C. Coverslips were peeled from mold, blocks were trimmed and sectioned. Ultrathin sections contrasted with 3% uranyl acetate for 5 min and 3% Reynolds lead citrate for 5 min. Samples were viewed using a JEOL JEM-1400 TEM at 100KV and digital images acquired with an AMT digital camera.

### Statistical analysis

The p values were calculated using a standard unpaired Student’s *t*-test with a p ≤0.05 considered significant. All statistical analyses were performed using at least three biological replicates.

## Acknowledgements

The plasmid expressing γ_1_ 34.5 - Flag was a gift from Dr. He Bin (University of Illinois, Chicago).

